# Out-of-equilibrium statistical dynamics of spatial pattern generating cellular automata

**DOI:** 10.1101/151050

**Authors:** Eduardo P. Olimpio, Hyun Youk

## Abstract

How living systems generate order from disorder is a fundamental question^1-5^. Metrics and ideas from physical systems have elucidated order-generating collective dynamics of mechanical, motile, and electrical living systems such as bird flocks and neuronal networks^6-8^. But suitable metrics and principles remain elusive for many networks of cells such as tissues that collectively generate spatial patterns via chemical signals, genetic circuits, and dynamics representable by cellular automata^1,9-11^. Here we reveal such principles through a statistical mechanics-type framework for cellular automata dynamics in which cells with ubiquitous genetic circuits generate spatial patterns by switching on and off each other’s genes with diffusing signalling molecules. Lattices of cells behave as particles stochastically rolling down a pseudo-energy landscape – defined by a spin glass-like Hamiltonian – that is shaped by “macrostate” functions and genetic circuits. Decreasing the pseudo-energy increases the spatial patterns’ orderliness. A new kinetic trapping mechanism – “pathway trapping” – yields metastable spatial patterns by preventing minimization of the particle’s pseudo-energy. Noise in cellular automata reduces the trapping, thus further increases the spatial order. We generalize our framework to lattices with multiple types of cells and signals. Our work shows that establishing statistical mechanics of computational algorithms can reveal collective dynamics of signal-processing in biological and physical networks.

---

A central question about living systems is how they generate ordered structures from disordered and thermally fluctuating mixtures of components^1-5^. Collective dynamics of interacting components that yield ordered states are often complicated but they arise from a few simple rules^1,9^. Striking examples of this include the spatial pattern formations in Conway’s “game of life”, collective migrations of birds^6,7^, and collective spikes in neuronal networks^8,10^. We can often explain collective dynamics of living systems composed of mechanical, motile, and electrical parts via theoretical frameworks such as the Vicsek^6,7^ and mechano-chemical models^2-4^ whose roots lie in statistical or soft matter physics. But such general frameworks are lacking for understanding the dynamics of an important class of systems – networks of cells such as tissues and biofilms in which multiple cells, without any morphogen gradients, collectively generate spatial patterns by regulating each other’s genes with diffusing chemicals and genetic circuits^9^. Indeed, it is unclear if and how principles such as information-optimization^12^ and metrics such as thermodynamic potentials, despite their utility in other contexts^13-19^, apply to spatiotemporal gene regulations by hundreds of cells. Moreover, we usually do not know which metrics suitably describe these systems. A core feature of these systems is that they are nonlinear dynamical systems that cellular automata can represent^1,9,11^ (Fig. 1a). To study them, we must often resort to exhaustive simulations that numerically solve a large number of chemical kinetics equations^9,11^. A wide-open question is if there are governing principles for these cellular automata dynamics that take us beyond the exhaustive simulations.

**Figure 1.**
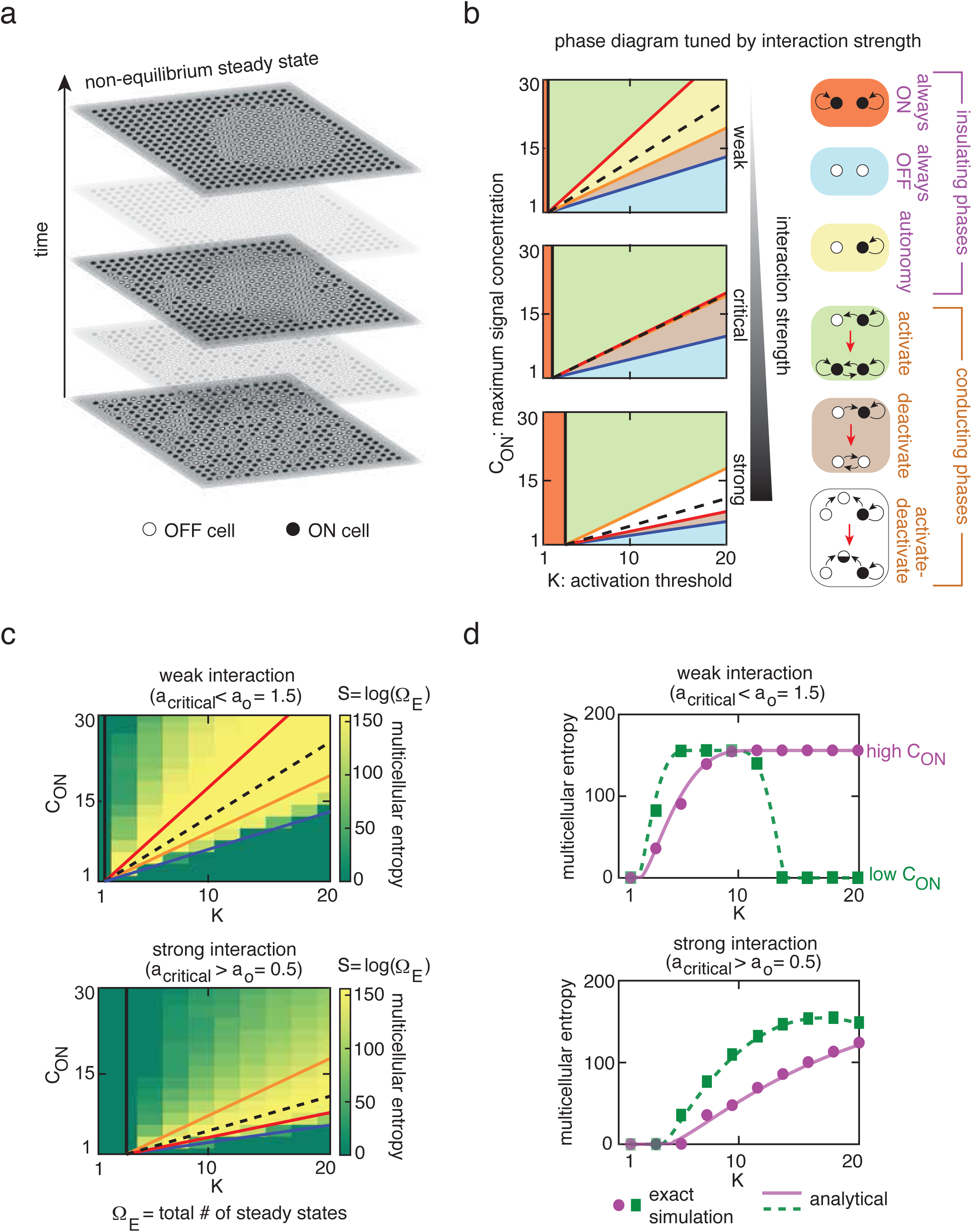
Steady states of the cellular automaton - Phase diagrams and multicellular entropy. **a,** Example of disorder-to-order transition driven by our cellular automaton. The top layer shows a steady state spatial pattern formed by ON-cells (black circles) and OFF-cells (white circles). **b,** Phase diagrams denoting all possible operational phases of the cellular automaton. Each colour represents a unique phase. Coloured lines represent phase boundaries that shift as the interaction strength is tuned. Insulating phases (orange, blue, yellow) allow only self-interaction while conducting phases (green, brown, white) allow cells to exchange information through their common signal such as ON-cell (black circle) activating an OFF-cell (white circle) into an ON-cell at the next time step (in the activate-phase). Black arrows represent interactions. Red arrows represent a flow in time. Phase diagram of the cellular automaton that operates with (top) a weak cell-cell interaction strength (*f*(*N*, *a_0_*) = 0.518; a_0_ = 1.5), (middle) a nearly critical cell-cell interaction strength (*f*(*N*, *a_0_*) = 0.967; a_0_ = 1.0), and (bottom) with a strong cell-cell interaction strength (*f*(*N*, *a_0_*) = 2.302; a_0_ = 0.5). **c,** Multicellular entropy, *S* = *log*(*Ω_E_*), computed for every phase and regimes of interaction strengths by running many cellular automata with *N*=121 (lattice of 11x11 cells), *R*=0.2*a_o_*, and a critical lattice constant *a_c_*~0.97 (where *f*(*N*, *a_c_*)=1). **d,** Analytical estimate of the multicellular entropy *S* (purple and green lines) compared with the exact values of *S* (purple and green data points) obtained for weak (top) and strong (bottom) interaction strengths, and the same parameters as in (c). Low *C_ON_* (*C_ON_* = 9.3) and high *C_ON_* (*C_ON_* =19.6). Each colour represents a horizontal slice (i.e. fixing *C_ON_* and varying K) taken in the two heat maps shown in (c).

Here we present a theoretical framework that, through new metrics and principles which are inspired by ideas from non-equilibrium statistical physics, explain and predict the dynamics of spatial pattern formation in a widely applicable and biologically relevant class of cellular automata. More generally, we present a framework for statistical dynamics of collective signal-processing by a network of living or non-living entities that digitally store information and interact through diffusing signals. We first consider cellular automata in which a finite number of identical cells on a lattice interact by exchanging a signaling molecule that flips on and off each other’s gene expression. We classify all spatial patterns of these “ON” and “OFF” autocrine cells^20,21^, which are ubiquitous in nature, into macrostates. The temporal evolution of the macrostates can be described by particles, which represent the cellular lattices, that flow along streamlines – akin to particles that drift and diffuse according to Langevin dynamics – in a phase space whose coordinates specify the spatial patterns’ orderliness. We find that the particles’ drifting-and-diffusing motions are in fact produced by a pseudo-energy landscape on which they stochastically roll down. We find that the pseudo-energy landscape’s shape is tuned by the strength of cell-cell communication, genetic circuits, and the macrostate functions of the cellular lattice. We show that a decrease in the particle’s (cellular lattice’s) pseudo-energy increases the orderliness of the spatial pattern and demonstrate a link between the pseudo-energy and spin glass’ Hamiltonians^22,23^. Unlike physical energies, however, we discover that the particles can be trapped at sloped regions of the pseudo-energy landscape which has no local minima and that this trapping is due to a new form of kinetic trapping – “pathway trapping” – that we uncover. The traps cause metastable spatial patterns. Noise in cellular automata can increase the spatial patterns’ orderliness by freeing the particles from the traps. Finally, we extend our framework to cellular automata with arbitrary numbers of cell types and of signal types. We find that a cellular lattice with multiple cell types behaves like a system of multiple interacting particles whose total pseudo-energy, which is the sum of each particle’s pseudo-energy and pseudo-interaction potentials, monotonically decreases over time. Our work suggests that establishing statistical mechanics of computational algorithms^24,25^, as we do here, is a promising way to understand how network-level dynamics of signal-processing emerge from a complex web of molecular and cellular interactions that are the basis of biological processes^6^. Moreover, since our statistical mechanics-type formalism also applies to non-living systems such as networks of signal-relaying switches, similar approaches may be fruitful in the future for understanding the network-level dynamics of signal-processing in physical networks.

Let us begin by formulating the rules for a cellular automaton that are inspired by multicellular systems such as the embryos of the fly *D. melanogaster*, hair follicles of mice, embryos of the frog *X. laevis*, zebrafish embryos, and microbial biofilms (see Supplementary Table 1 for details of these systems). We consider a two-dimensional lattice of *N* identical, spherical cells of radius *R* that communicate via “autocrine” signalling^20,21^ (Supplementary Fig. 1). Autocrine signalling, along with paracrine signalling that we will later consider, accounts for a vast majority of cellular communications in nature^20^. Each autocrine cell in the lattice interacts with itself and all the other cells by secreting a signalling molecule (“signal”) that diffuses outside the cells (details in Supplementary section 1.1). Each cell senses the concentration *c* of the signal on its surface. This in turn determines each cell’s signal-secretion rate and the expression level of its “output gene”. We can consider lattices of any shape but for concreteness, we focus here on a triangular lattice of cells with a lattice spacing *a_o_*. Motivated by experimental findings^15,20,21^, we consider autocrine cells with a switch-like response to the signal (Supplementary section 1.1). Namely, if the value of *c* on the cell is larger than a threshold concentration *K*, then the cell switches “ON” its output gene. Otherwise (i.e., *c* < *K*), the cell switches “OFF” its output gene. The ON-cell secretes the signal at a constant rate such that a lone ON-cell (i.e., *N*=1) would maintain a signal concentration of *C_ON_* on itself whereas a lone OFF-cell would constantly secrete the signal at a lower rate such that it would maintain a signal concentration of 1 on itself (*C_ON_* > 1). Each cell can thus interact with itself through its own signal (i.e., a cell engages in “self-interaction”). We assume that the cells take longer times to adjust their ON/OFF states than for the diffusing signal to reach steady state concentrations across the lattice (see Supplementary section 1.1. for justifications). These rules define a cellular automaton that proceeds as follows: (1) compute the steady state value of *c* on every cell, (2) simultaneously update each cell’s state according to the value of *c* on each cell, and (3) go back to step (1). The cellular automaton terminates when no cell’s state requires further updates (details in Supplementary section 1.1). The terminal spatial patterns formed by the ON/OFF cells are out-of-equilibrium steady states that are sustained as long as there is a continuous input of energy to fuel the cells’ signal secretions^1^. Intriguingly, we found that this cellular automaton could drive disordered spatial patterns into many highly ordered steady state spatial patterns (Fig. 1a). We sought to reveal the principles that drive these pattern formations.

Let us begin our analysis of the cellular automaton by noting that we can classify the cellular automaton into distinct operational “phases”. Just as the phases of matter such as the gas phase and the solid phase characterize the interactions among the constituent particles, the phase of the cellular automaton characterises the type of cell-cell interaction that drives the cellular automaton. From the start to the end of a cellular automaton, its phase remains fixed and is determined by the values of *K*, *C_ON_*, and an “interaction strength” *f_N_*(*a_o_*) that are held constant over time. The interaction strength quantifies the connectivity among all cells formed by the diffusing signal and is defined as 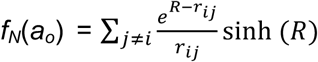, where *r_ij_* is the distance between cell-i and cell-j. Using methods that we established in a previous work^26^ (details in Supplementary section 1.2), we obtained phase diagrams for the cellular automaton as a function of *K*, *C_ON_*, and the interaction strength (Fig. 1b). All phases belong to either an “insulating phase” or a “conducting phase”. Varying the interaction strength tunes the transitions between the insulating phases – in which there is no information exchange among cells because each cell is inert to the other cells’ signal (i.e. a cell cannot influence any other cell) – and “conducting” phases – in which the cells exchange information with others through their signal (i.e. a cell can influence the other cells) (Fig. 1b and Supplementary Fig. 2). When the interaction is weak (i.e., *f_N_*(*a_o_*) < 1), the cellular automaton can operate in the “autonomy” phase – an insulating phase in which each cell, due to its self-interaction, can be stably ON or stably OFF regardless of the other cells’ states. Also, weak interactions (*f_N_*(*a_o_*) < 1) enable the cellular automaton to operate in conducting phases such as the “activate” phase – in which enough neighbouring ON-cells can turn on an OFF-cell – and the “deactivate” phase – in which enough neighbouring OFF-cells can turn off an ON-cell. At a critical interaction strength (i.e., *f_N_*(*a_o_*) = 1), the autonomy phase disappears and the boundaries of the activate and deactivate phases start to merge (Fig. 1b and Supplementary Fig. 2). When the interaction is strong (i.e., *f_N_*(*a_o_*) > 1), an “activate-deactivate” phase appears in which both activation and deactivation can occur (Fig. 1b). We found that the cellular automaton in this phase could yield some of the most highly ordered steady state spatial patterns such as islands and stripes of ON-cells (Fig. 1a).

The phase diagrams enable us to organize our analysis of the cellular automaton’s dynamics. The phase of an automaton characterizes the type of cell-cell interaction that drives the spatial pattern formation dynamics. We can ask how each type of cell-cell interaction (i.e., each phase) determines the disorder-to-order dynamics (Fig. 1a). To address this question, we began by asking what the total number *Ω_E_* of steady state spatial patterns that can be produced in each phase is. There are 2^N^ possible patterns that the cellular automaton can start with. For the cellular automaton that operates in the autonomy phase, all of the 2^N^ patterns are steady states because each cell is perfectly insulated from the others’ signal. But in the conducting phases, cell-cell interactions would constrain where each ON- and OFF-cells can be placed on the lattice to form steady state patterns. By running many cellular automata, we obtained the exact value of “multicellular entropy”, *S* = *log*(*Ω_E_*), for each phase (Fig. 1c and Supplementary Fig. 3; details in Supplementary section 1.3). We found that increasing the interaction strength tends to decrease the multicellular entropy, suggesting that the cells’ states become more correlated (Fig. 1c). Interestingly, we found that in the strong interaction regime, the multicellular entropy for the activate-deactivate phase – a conducting phase – can be almost as high as it is in the autonomy phase (Fig. 1c – between orange and red line in “strong interaction”) as if to suggest that conduction of signal among cells no longer matters. We will later see that this is due to a form of kinetic trapping that is hidden from our view at the moment. To better understand the property of the multicellular entropy, we derived an analytical estimate of *S* as a function of *C_ON_*, *K*, and *f_N_*(*a_o_*) by improving a method that we introduced in a previous work^26^ (details in Supplementary section 1.3). Our analytical estimate of the multicellular entropy closely matches the exact values of *S* obtained from the simulations (Fig. 1d). The multicellular entropy gives a qualitative sense of how long it takes for the cellular automaton to terminate. Intuitively, we expect that the cellular automaton that operates in a phase with a high value of *S* may terminate relatively quickly because there is a high chance that there exists a steady state spatial pattern “close to” its initial spatial pattern. This is certainly true in the autonomy phase, in which every cellular automaton halts in zero time steps because every pattern is a steady state. To test if our intuition applies to the other phases, we have to more deeply understand the dynamics of the cellular automaton. We turn to this issue next.

So far, we have classified the cellular automaton into distinct phases and determined the number of steady states in each phase. Let us now look for principles that drive the dynamics that lead to the steady state patterns. Running the cellular automaton requires keeping track of every cell’s state and the signal concentration on every cell for each time step^11^. Since the number of parameters increases exponentially as the number *N* of cells increases, analytically predicting (i.e., without running the cellular automaton) the exact temporal evolution of any one spatial pattern is infeasible even for populations of a modest size (e.g., *N* ~ 100). But just as we can describe a box of gas with macrostate variables such as temperature without specifying every gas particle’s position and momentum, we sought to define macrostates of the cellular lattice – which represent ensembles of specific spatial patterns (“microstates”). We explored two macrostate functions (Fig 2a) – the fraction *p* of the *N* cells that are ON, and a “spatial order parameter” *I* that we define as

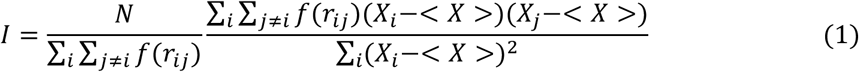

Where 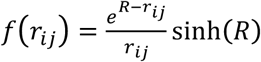 is the interaction term for the cell pair (*i*, *j*) that appears in the interaction strength function *f_N_*(*a_o_*), *X_i_* is +1 for an ON cell-i and −1 for an OFF cell-i, and 〈·〉 is the average over all cells (see examples of spatial patterns that correspond to each value of *I* in Fig. 2a and Supplementary Fig. 4). The spatial order parameter assigns a single value to each spatial pattern. It measures the auto-correlation of the cell states that is weighted by the strength of each cell pair’s interaction (details in Supplementary section 2.1). The magnitude of the spatial order parameter |*I*|, lies between zero and one. As |*I*| approaches 0, the spatial pattern becomes more disordered. As |*I*| approaches 1, the spatial pattern becomes more ordered. With *p* and *I* as macrostate functions, we may view a cellular lattice as a particle that moves in a phase space whose coordinates are *p* and *I*. Since many spatial patterns (“microstates”) can belong to the same macrostate (*p*, *I*) (Fig. 2a) and the cellular automaton, a priori, seems to care only about microstates rather than macrostates, multiple particles with the same initial position (*p*, *I*) might move in highly divergent paths in the phase space. Thus, it is unclear that a macrostate-level description, such as an equation of motion (*p*(*t*), *I*(*t*)) as a function of time *t*, for spatial pattern formation is even feasible.

**Figure 2.**
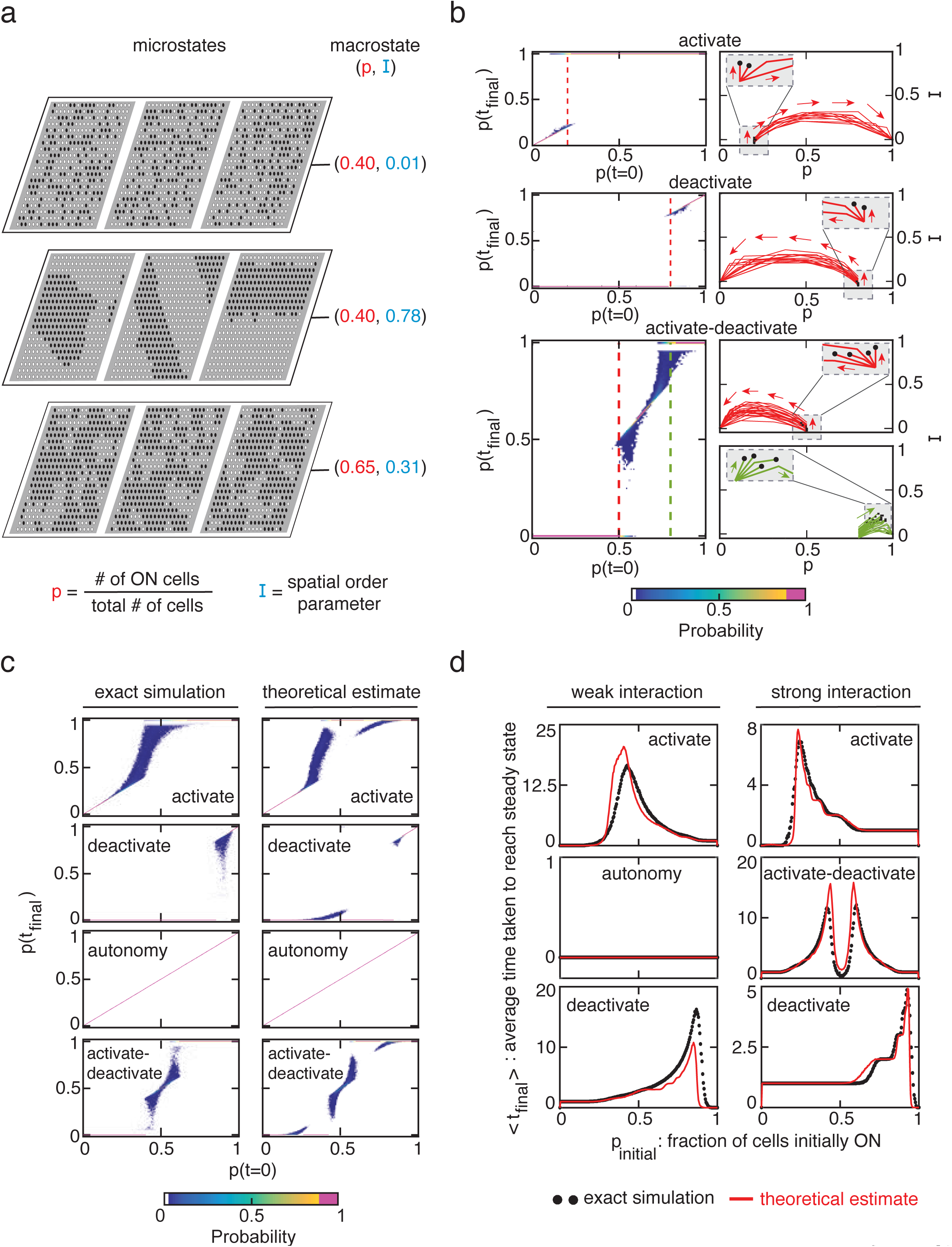
Macrostate description - Cellular lattices act as particles that drift and diffuse in a phase space of spatial patterns. **a,** A macrostate (*p*, *I*) of cellular lattice (right column) is an ensemble of microstates (i.e., individual patterns) of cellular lattice that have the same *p* and *I*. **b,** (Left column) Probability density maps showing *p*(*t_final_*) - the steady state value of *p* when the cellular automaton terminates at time *t_final_* - as a function of *p*(*t*=0) - the initial value that the automaton starts with. For each value of *p*(*t*=0), we ran at least 1000 cellular automata, each with the same parameters but with a different, completely disordered microstate (*I* ≈ 0) that we randomly chose. Each coloured “pixel” here represents a probability (see colour bar at the bottom). White regions represent values near zero probability. Dashed red and green lines are slices along the density maps that correspond to the trajectories in the right column. (Right column) Macrostate trajectories (*p*(*t*), *I*(*t*)) as a function time *t*. For each phase, 30 trajectories that start with the same macrostate are shown (red curves). Arrows represent general directions of flow in each trajectory. Insets in each panel show zoomed-in views of a few trajectories. Black dots in the insets represent termination of trajectories. In each of the three conducting phases, all 30 trajectories start with the same initial values of *p* and *I* - Activate phase: *p*(*t*=0)=0.2 (with *K*=14, *C_ON_*=16); deactivate phase: *p*(*t*=0)=0.8 (with *K*=18, *C_ON_*=7); activate-deactivate phase: *p*(*t*=0)=0.5 for red trajectories and *p*(*t*=0)=0.8 for green trajectories (with *K*=16, *C_ON_*=8). For all three phases, *N*=121 and *R*=0.2*a_o_*. **c,** (Left column) Probability density maps obtained as in (b). (Right column) With the same parameters as in the left column, probability density maps obtained by Monte Carlo simulations based on our theory of drifting-and-diffusing particles (with analytical estimates of key probability distributions) and our “branching algorithm” (see Supplementary sections 2.3 and 2.4). Activate phase: *K*=20, *C_ON_*=15, *a_O_*=1.5; deactivate phase: *K*=6, *C_ON_*=15, *a_O_*=1.5; autonomy phase: *K*=12, *C_ON_*=15, *a_O_*=1.5; Activate-deactivate phase: *K*=15, *C_ON_*=8, *a_O_*=0.5. In all phases, *N*=225 and *R*=0.2*a_O_*. **d,** Average number < *t_final_* > of time steps taken by the cellular automaton to terminate as a function of the initial value of p, *p_initial_*. Each black data point is an average over at least 1000 cellular automata runs for each value of *p_initial_*. Red curves are analytical estimates based on our theory of drifting-and-diffusing particles in phase space.

To explore the feasibility of obtaining a macrostate-level description of an initially disordered pattern (*p*(*t* = 0) = *p*_initial_, *I*(*t* = 0) = 0) evolving into a steady state spatial pattern (*p*(*t*_final_), *I*(*t*_final_)) after *t*_final_ time steps, we ran the cellular automaton for all possible values of *p*_initial_. Specifically, for each value of *p*_initial_, we randomly chose hundreds of microstates that belong to the same macrostate (*p*_initial_, 0) and then ran the cellular automaton (details in Supplementary section 2.2). In doing so, we obtained, for each phase of the cellular automaton, a distribution of values for *p*(*t*_final_) as a function of *p*_initial_ (Fig. 2b and Fig. 2c – left columns). The fact that, for a given value of *p*_initial_, we obtain a distribution of rather than a single value of *p*(*t*_final_), shows that if a macrostate-level description exists, then it would need to be stochastic. We also obtained an ensemble of possible trajectories (*p*(*t*), *I*(*t*)) of the particles in the phase space (Fig. 2b – right column; Supplementary Figures 5-7). Intriguingly, we found that the particles that start at the same position (*p*_initial_, 0), for the most part, remained close to each other in subsequent times, which led to tightly bundled streamlines in the phase space (Fig. 2b – right column). Moreover, in all the trajectories, we find that the *I* first increases and then plateaus while the *p* either increases or decreases over time (Fig. 2b). Then, after some more time passes, we observed one of two events occurring. In some of the trajectories, the *I* decreases to zero while the *p* either approaches zero or one. In the other trajectories, the particles stop at non-zero values of *I* and *p*. The later scenario occurs most notably but not exclusively in the activate-deactivate phase (Fig. 2b – insets show black dots terminating trajectories). Taken together, the regularities that we observed in the particle trajectories suggested that an equation of motion, which is stochastic, might underlie the dynamics of the particles in the phase space – an idea that we next turn to.

Given the particles’ stochastic flows in the phase space, finding equations of their motion reduces to determining the probability *P*(*p*(*t+1*), *I*(*t+1*) | *p(t)*, *I(t)*) that a particle at position (*p(t)*, *I(t)*) at time *t* moves to position (*p(t+1)*, *I(t+1)*) at the next time step. We can rewrite equation (1) as

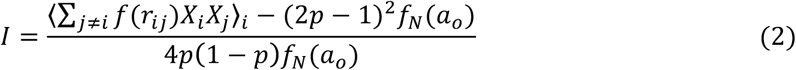

where 〈·〉_*i*_ is the average over all *N* cells. Equation (2) shows that the *I* depends on *p* and suggests that the particles’ trajectories should be constrained, as we have indeed previously observed (Fig. 2b – right column). Equation (2) also shows that as *p* approaches zero or one, *I* suddenly “collapses” to zero, as we also observed in the simulations (Fig. 2b – right column; see Methods and Supplementary section 2.1). We hypothesized that the particles’ stochastic flows result from a drift-diffusion process, with some “driving force” in the cellular automaton causing the cellular lattice to drift towards more ordered patterns while our ignorance about which microstate represents the spatial pattern at each time acting as a source of diffusion. Specifically, by denoting the 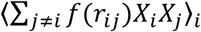 term in equation (2) as Θ, which we call the “normalized spatial order”, our idea would suggest that *p* and Θ would follow

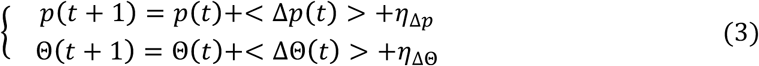

where Δ*p*(*t*) ≡< *p*(*t* + 1) – *p*(*t*) >, ΔΘ(*t*) ≡< Θ(*t* + 1) – Θ(*t*) >, *η*_Δ*p*_ is the noise in *Δp*, and *η*_ΔΘ_ is the noise in *ΔΘ*. The “noise” terms here represent our ignorance of the microstates – we do not know the precise location of every ON- and OFF-cells on the lattice at any given time – rather than representing biological sources noise such as noise in gene expression (we will turn to biological noise later). Since *p* and Θ uniquely determine *I*, equation (3) specifies *I*(*t*) as well. Indeed we could determine the probability *p*(y_-_, y_+_; n) that y_-_ ON-cells turn off and y_+_ OFF-cells turn on in the next time step, given that there are *n* ON-cells (details in Supplementary section 2.3). Furthermore, we could determine the means and the variances of Δ*p*(*t*) and ΔΘ(*t*) (details in Supplementary section 2.4). In turn, we could determine how a given macrostate (*p*(*t*), Θ(*t*)) stochastically evolves over time through equation (3). To test how well the equation of motion describes the particle trajectories, we must use Monte Carlo simulations that are based on our analytically determined statistical properties of Δ*p*(*t*) and ΔΘ(*t*) because equation (3) is inherently stochastic. We developed a new algorithm – “branching algorithm” – for performing such Monte Carlo simulations. The algorithm has similarities to the “pruning algorithm” for simulating two-player games and protein folding (Supplementary Figs. 8-9; details in Methods and Supplementary Sections 2.3 and 2.5). Comparing the particle trajectories from our Monte Carlo simulations with those of the cellular automaton, we found that the main statistical features of the trajectories produced by the two different methods closely agreed (Figs. 2c and 2d; Supplementary Figs. 10-14). Namely, the Monte Carlo simulations closely estimate the distribution of values of *p*(*t*_final_) for each value of *p*_initial_ (Fig. 2c) as well as the mean termination times 〈*t*_final_〉 of the cellular automaton (Fig. 2d). Thus, the cellular lattice indeed acts like a particle that drifts and diffuses in the phase space. Crucially, we find that the particle’s drift causes it to increase its value of Θ over time, which means that the spatial pattern that the particle represents becomes more ordered over time. While the diffusive motion is due to the probabilistic description that arises from our ignorance of the precise microstates of the cellular lattice at each time, we have not yet addressed what the supposed driving force that causes the particles to drift towards higher values Θ is. We turn to this issue next.

Let us now understand what causes the drift that is responsible for the normalized spatial order monotonically increasing over time (Fig. 2b – insets in the right column). To address this, we adapted the technique used in analysing the 2D Ising model’s critical phenomena. Suppose that we enclose islands of ON-cells by polygons (Fig. 3a). By averaging over all ON-cells, we find that the mean number < *m_ON_* > of ON nearest neighbours of any cell is 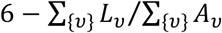, where {*v*} is the set of all polygons, *L_v_* is the number of edges for polygon-*v*, and *A_v_* is the number of ON-cells inside polygon-*v*. By developing a mean-field approach, in which we analyze how the polygonal islands grow over time and compete in a field of randomly distributed ON- and OFF-cells (details in Supplementary section 3.1), we found that both the < *m_ON_* > and the total area of the polygons, 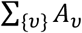, monotonically increase over time (Supplementary Figs. 15-17). The growing and the merging of the polygonal islands of ON-cells over time is synonymous with the spatial order parameter increasing over time until it potentially collapses (as *p* approaches 0 or 1). More concretely, with the mean-field approach, we could estimate how the spatial order parameter changes over time in terms of < *m_ON_*(*t*) > and *p*(*t*) (details in Supplementary section 3.2):

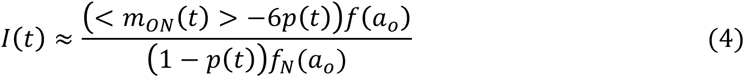

**Figure 3.**
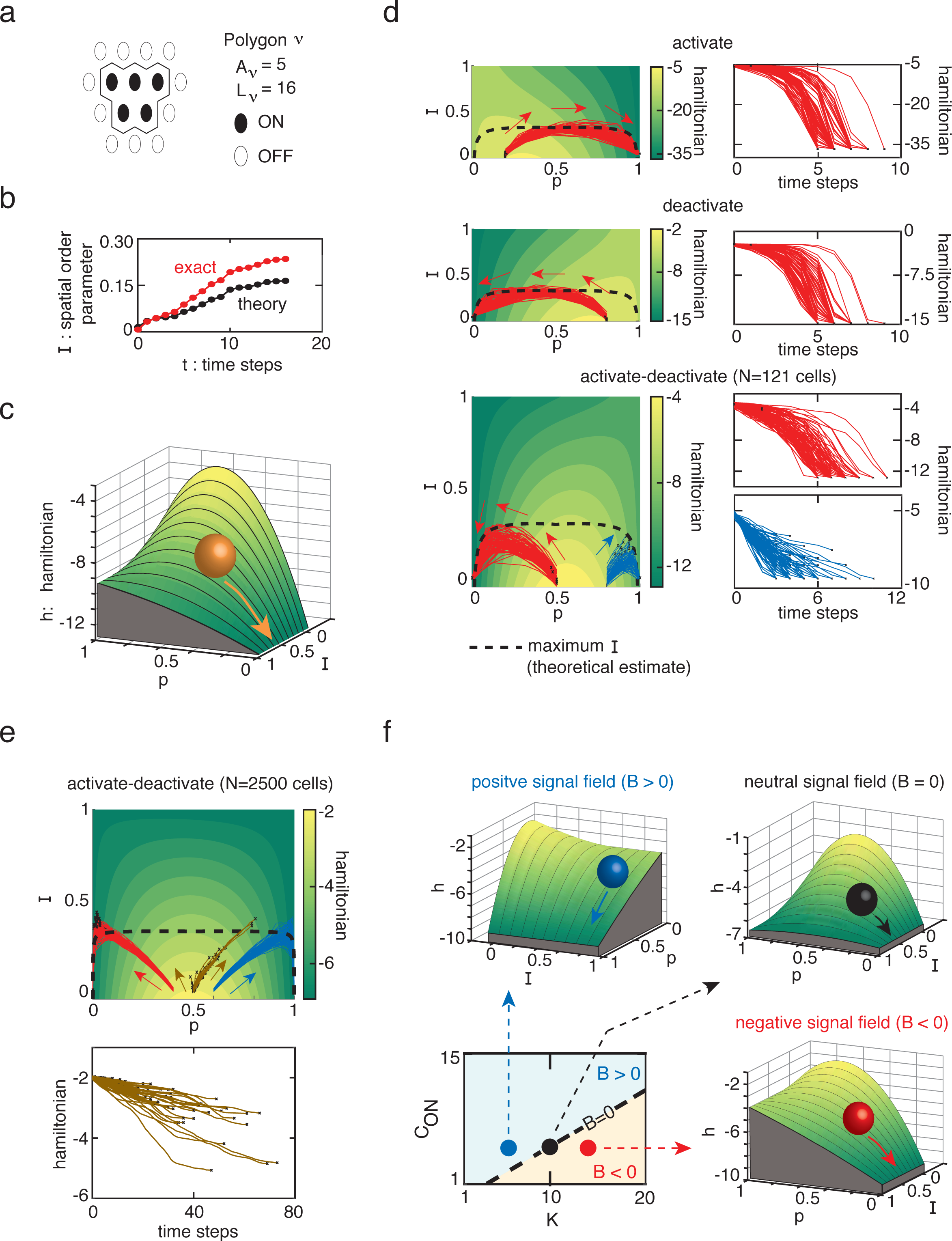
Cellular lattices act as particles that drift and diffuse downwards and get stuck along inclines of pseudo-energy landscapes. **a,** Example of the polygonal analysis, with polygon-*v*, for studying disorder-to-order transition. *L_v_* and *A_v_* are the number of edges and the number of enclosed ON-cells for polygon-*v*. **b,** Comparing the analytical formula for *I*(*t*) (equaton (4) - black curve) obtained by the polygonal analysis shown in (a) with the exact value of *I*(*t*) obtained from a cellular automaton (*N*=121, *a_O_*=0.5, *R*=0.2*a_O_*, *K*=10, *C_ON_*=5). **c,** Pseudo-energy landscape obtained by plotting *h*(*p*, *I*) = *H/N* - multicellular Hamiltonian *H* divided by *N*. *N*=121, *a_O_*=0.5, *R* = 0.2*a_O_*, *K*=16, *C_ON_*=8. Particle (orange circle), representing a cellular lattice, rolls down the landscape. **d,** (Left column) Each phase shows 100 particle trajectories (red and blue curves), some of which appeared in Fig. 2(b), with the same parameters as in Fig. 2b but now with the pseudoenergy landscapes in the backgrounds (heat maps). Arrows indicate general directions of the particle flows over time. Black curves indicate theoretically predicted maximum possible value of *I* as a function of *p* (see Supplementary section 3.2). (Right column) Normalized Hamiltonian (*h* = *H/N*) for each trajectory in the left column. All are for *N*=121 cells. **e,** (Top) Same parameters as the activate-deactivate phase in (d) but now with more cells (*N*=2500). Red trajectories start with *p*=0.4, brown trajectories start with *p*=0.5, and blue trajectories start with *p*=0.6. Black curve indicates theoretically predicted maximum possible value of *I* as a function of p. (Bottom) Normalized Hamiltonian (*h* = *H/N*) for each brown trajectory in top panel. **f,** Pseudo-energy landscapes for three different regimes of the signal field *B* in and that boundaries of the activate-deactivate phase: (Top left) At a boundary between activate and activate-deactivate phases (*B*=2.6, *K*=7.3, *C_ON_*=5), (Top right) Within activate-deactivate phase (*B*=0, *K*= 9.9, *C_ON_*=5), (Bottom right) At a boundary between deactivate and activate-deactivate phases (*B*= −2.6, *K*=12.5, *C_ON_*=5). (Bottom left) Corresponding phase diagram denoting signs of *B*. In all cases, *N*=225, *a_O_*=0.5, *R*=0.2*a_O_*.

We found that equation (4) closely agrees with the exact value of *I*(*t*) obtained from cellular automata that start with highly disordered (*I* ~ 0) spatial patterns (Fig. 3b). Moreover, equation (4) shows that if an initial pattern is highly disordered (*I* ~ 0), then we have <*m_ON_*(*t* = 0)> = 6p(*t* = 0) and that in subsequent times, we have < *m_ON_*(*t*) > ≥ 6*p*(*t*) because the < *m_ON_* > monotonically increases over time. In other words, the *I*(*t*) monotonically increases over time. In addition, equation (4) predicts that the *I* has a maximum for each value of *p* (details in Supplementary section 3.2). This maximum spatial order parameter *I_max_* (Fig. 3d – black curves) closely matches the actual maximum reached by the particle trajectories in the cellular automata (Fig. 3d – red curves). In summary, we find that the disorder-to-order transition is associated with islands of ON- and OFF-cells competing for growth and this competition leads to maximally allowed spatial orders.

So far, we have established that the cellular lattice behaves as a particle that drifts and diffuses in the phase space and that we can understand the spatial order parameter monotonically increasing over time by focusing on competition between islands of ON- and OFF-cells. A missing piece in our understanding is the mechanism that drives the temporal changes in *p*, and in turn, perhaps a deeper reason behind why the spatial order parameter tends to increase over time (since *I* depends on *p*). It is unclear, so far, what determines whether *p* increases or decreases over time in each phase. For example, in the activate phase, some values of *p*_initial_ cause more cells to turn on over time while for others, *p* does not change (Fig. 2c). Similarly, in the activate-deactivate phase, there appears to be a critical value of *p*_initial_ above which more cells turn on over time and below which more cells turn off over time (Figs. 2b and 2c). We found that we could explain these observations by defining a multicellular Hamiltonian *H*,

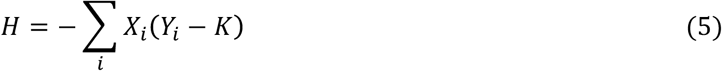

where *Y_i_* is the signal concentration on cell-i. We can rewrite the *H* in terms of *p* and *I* to obtain a nearly intrinsic function *h*(*p*, *I*) = *H*(*p*, *I*)/*N* (details in Supplementary section 3.3). By plotting *h* as functions of *p* and *I*, we obtain a “pseudo-energy” landscape (Fig. 3c) whose shape is determined by the phases of the cellular automaton (Fig. 3d). Importantly, we could prove that *h* is a Lyapunov function – *h* decreases or stays the same over time until the cellular automaton halts (proof in Supplementary section 3.3). Indeed, reminiscent of potential energy landscapes of physical particles, plotting the trajectories of our pseudo particles (same as in Fig. 2b) on top of their respective pseudo-energy landscapes revealed that the particles stochastically “roll” down (i.e., drift-and-diffuse down) their pseudo-energy landscapes until they halt (Fig. 3d). The location of the pseudo-energy landscape’s peak, which occurs at *I*=0 for all three conducting phases (Fig. 3d), determines for which values of *p*_initial_ the particle drifts towards increasing values of *p* in subsequent times. But unlike physical energy landscapes whose local minima trap physical particles, the pseudo-energy landscape lacks any local minima and traps many particles at its sloped regions. This trapping is particularly but not exclusively evident in the activate-deactivate phase (Fig. 3d – blue curves; Fig. 3e – brown curves). We will address what causes this trapping after addressing what determines the shape of the pseudo-energy landscape.

Let us explore how one can tune the shape of the pseudo-energy landscape, and in the process, show that the multicellular Hamiltonian is analogous to, but not the same as, the Hamiltonians of spin glass^22,23^ and the Hopfield network^27,28^. We will then show that the trapping mechanism is similar in spirit to, but not the same as, the notion of frustration in spin glasses. In the activate-deactivate phase, increasing the number *N* of cells causes the pseudo-energy landscape to be more “left-right” symmetric (i.e., more symmetric about the line that joins (*p* = *p*_peak_, *I* = 0) and (*p* = *p*_peak_, *I* = 1), where *p*_peak_ is the location of the landscape’s peak) (i.e., compare the activate-deactivate phase in Fig. 3d with Fig. 3e). Making the landscape more symmetric enables more particles that start at a position near (*p*_peak_, *I* ~ 0) to bifurcate: some particles, according to the equation of motion, have a probability of moving down the pseudo-energy landscape towards decreasing values of *p* (i.e., Δ*p* < 0) while others have a probability of moving down the pseudo-energy landscape towards increasing values of *p* (i.e., Δ*p* > 0) due to diffusion (Fig. 3e – brown paths). Taking the “thermodynamic limit” (i.e., *N* → ∞) in the activate-deactivate phase, we found that the landscape becomes perfectly symmetric and that the equations of motion (equation (3)) can now predict particle trajectories for cellular automata that are infeasible to run due to the large value of *N* (details in Supplementary section 3.3). To more systematically under the landscape’s shape, we can rewrite *H* as

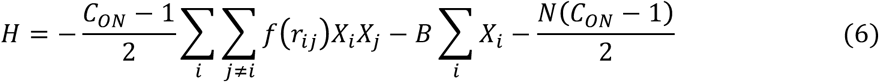

where *B* is a “signal field” defined as 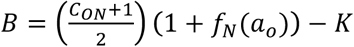. Equation (6) shows that the multicellular Hamiltonian is strikingly similar, at least in its form, to the Hamiltonian 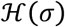 of the Hopfield network^27,28^ and magnetic spins with arbitrarily long range interactions (e.g., spin glass^22,23^), both of which take the form

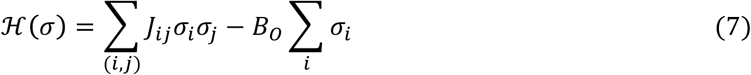

where *B_O_* is an external magnetic field or its equivalent, the first summation runs over every pair of spins or neurons *σ_i_* and *σ_j_* including those that are far apart, *J_ij_* and is a coupling constant for the spin or neuronal pair (*i*, *j*). But we stress that unlike the Hopfield network and spin glasses^22,23,27,28^, the pseudo-energy landscape has no local minima and that our particles can stop at inclined parts of the pseudo-energy landscape. Nonetheless, equation (6) tells us that the signal field *B* is a useful knob that one can tune to change the pseudo-energy landscape (Fig. 3f) and that it can compete with the cell-cell interaction term (with coupling constant 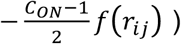 just as the magnetic field *B_O_* does in spin glasses and the Hopfield network^22,23,27,28^. Namely, a positive *B* corresponds to the activate phase, a negative *B* corresponds to the deactivate phase, and a zero *B* corresponds to the activate-deactivate phase (Fig. 3f; Supplementary Fig. 18; Supplementary section 3.3).

Having determined how the pseudo-energy landscape’s shape can be tuned, we now turn to explaining why particles are trapped at sloped regions of the pseudo-energy landscape. Due to the finite number of cells, the phase space – spanned by *p* and *I* – consists of a discretized “movement grid”: the *p* and *I* of a particle can only change in discrete units. Moreover, a particle at any one location in the phase space does not have the freedom to drift and diffuse in all directions because the *I* depends on the *p* (equation (2)). Thus, the movement grid consists of a complex web of alleys, turns, and dead-ends that can only be revealed by a microstate-level description – one that keeps track of the exact location of each ON- and OFF-cell at every time step. Thus, a particle may be located on the movement grid where it could decrease its pseudo-energy further by taking additional steps in a certain direction. But it may not be able to take those steps because no accessible paths exist in that direction and thus the particle is stuck at a dead-end. We can make this notion – which we call “pathway trapping” – more rigorous by defining “trapping configurations” that are similar in flavour to, but not the same as, the notion of frustration in spin glasses and idea of kinetic trapping. To find the trapping configurations, we can use the nearest neighbour approach that we have used for estimating the *I*(*t*) (equation (4)) (see details in Supplementary section 3.3). As an example, consider a cellular automaton, operating in the activate-deactivate phase, that generates a spatial pattern composed of a single island, call it “island-*v*”, of ON-cells with *A_v_* = 5 and *L_v_* = 16 (depicted in Fig. 3a). For this cellular automaton, we find that an OFF-cell, in order to turn on, requires at least three of its nearest neighbours be on while an ON-cell, in order to turn off, requires at least five of its nearest neighbours to be on. These two conditions cannot be simultaneously satisfied by the spatial pattern consisting of the island-*v* (Fig. 3a) despite the fact that turning more cells on (and thus simultaneously turning more cells off) would decrease the multicellular Hamiltonian. This is an example of a trapping configuration. As the *I* increases over time, the cellular automaton that operates in the conducting phases can realize such “trapping” conditions more and more. We find that the largest number of trapping configurations occur in the activate-deactivate phase. As a result, the activate-deactivate phase exhibits the highest multicellular entropy among all the conducting phases and allows for many metastable patterns, as we previously observed (Fig. 1c).

Until now, we have analysed deterministic cellular automata. But real cells are noisy – their gene expressions are stochastic^1,8-10,12-16,18,19^. We can treat such biological noise by having the secretion rate and the threshold *K* to fluctuate over time for each cell. The combined effect of fluctuating both *K* and *C_ON_*, however, would be equivalent to letting the *K* alone fluctuate for each cell (Fig. 4a). We have modified our cellular automaton so that at each time step and for each cell, we pick a new value of the threshold *K* + δ*K*, where *K* is the same constant for all cells and δ*K* is a normally distributed noise with a mean of zero and a variance of α^2^ (Fig. 4a; details in Supplementary section 4.1). We can then define a “noise strength”, *ξ* = *α*/*K*, which helps us determine how much noise is required to cause the particles’ trajectories in the phase space to deviate significantly from those of the deterministic cellular automata. Intuitively, we expect such deviations to occur if *α* is comparable to or larger than either |〈*Y_i_*_=*ON*_〉 − *K*| or |〈*Y_i_*_=*OFF*_〉− *K*|, where <*Y*_*i*=*ON*_> and <*Y_i_*_=*OFF*_> are the average signal concentrations on ON-cells and OFF-cells respectively. Indeed, we find that the minimum noise strength *ξ_min_* required to significantly perturb the deterministic cellular automata dynamics is 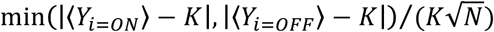 (Supplementary Fig. 19; details in Supplementary section 4.2). To confirm this, we ran cellular automata with low noise strengths (*ξ* < *ξ_min_*) and found that they did not yield *p*(*t*), *I*(*t*), and *h*(*t*) that were much different from those of the deterministic cellular automata. Cellular automata with high noise strengths (*ξ* > *ξ_min_*) indeed produced particle trajectories that differed from those of the deterministic automata. In particular, noise of a low strength cannot destroy a spatial pattern that is in a steady state of the deterministic cellular automaton (Fig. 4b – left column; Supplementary Fig. 20) while a noise of a high strength can drive a pattern out of a steady state by causing the *p* change, the *h* to decrease further, and the *I* to increase further until *p* reaches either zero or one (Fig. 4b – right column). Thus, the noise strength defines the stability of steady state spatial patterns.

**Figure 4.**
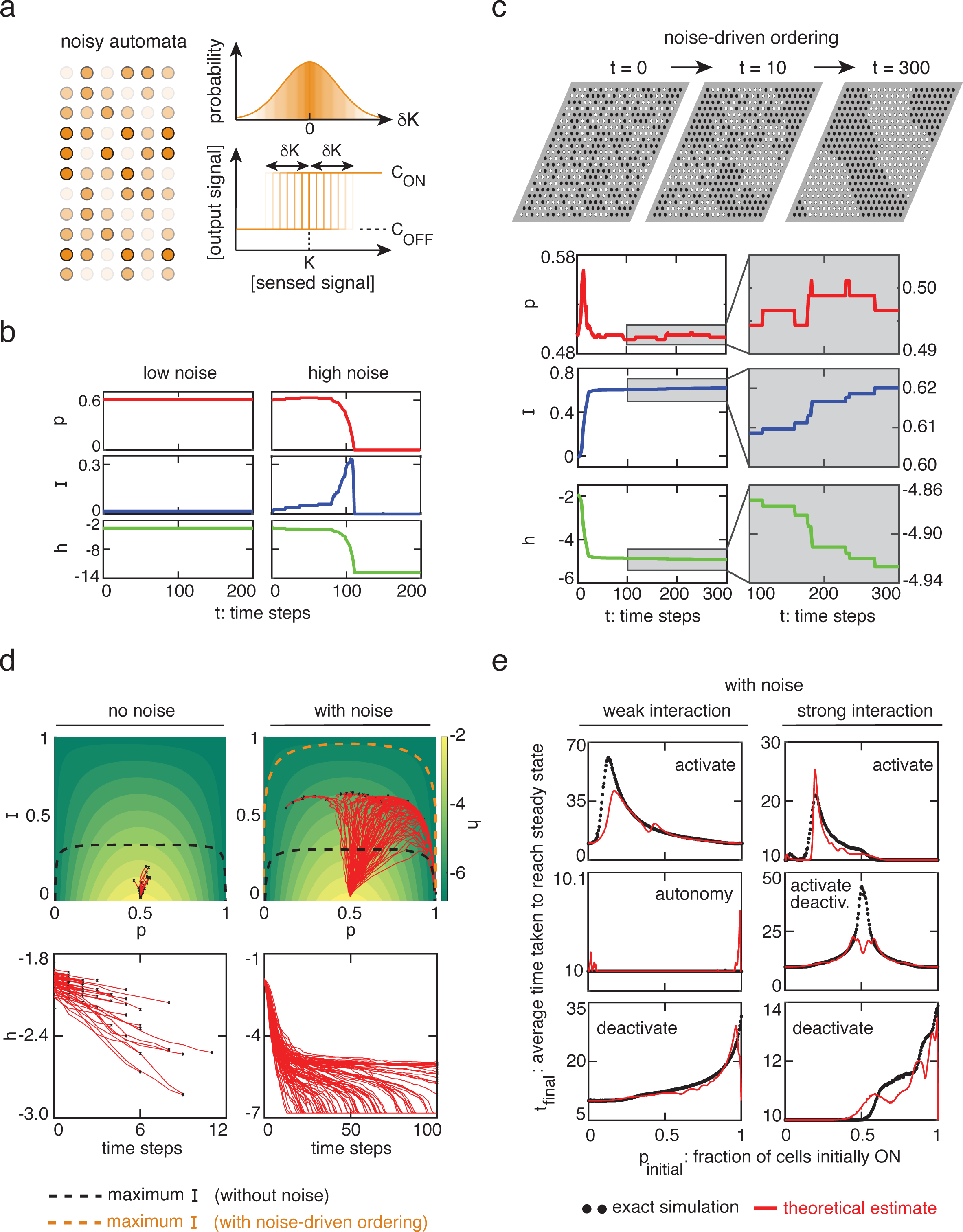
Stochastic cellular automata - Reducing pathway trapping creates metastable patterns with higher spatial orders. **a,** Schematics of noisy cellular automata. Each cell can have a different activation threshold *K* + δ*K*, where δ*K* is a Gaussian noise with a zero mean and variance *α*^2^ that remains fixed throughout the cellular automaton run. **b,** Examples of *p*, *I*, and *h* (=*H/N*) for low noise (α = 0.2) and high noise (α = 0.5). In both cases, *N*=121, *R*=0.2*a_O_*, *a_O_*=0.5, *K*=16, and *C_ON_*=8 (corresponds to activate-deactivate phase). Theoretically predicted minimum value of *α* required for perturbing deterministic automata dynamics here is 0.4 (see Supplementary section 4.2). **c,** Example of noise-driven disorder-to-order transition an activate-deactivate phase (*K*=10, *C_ON_*=5) with *p*(*t*=0)=0.50. Black circles are ON-cells and white circles are OFF-cells. Noise strength *ξ* =0.5, *N*=441, *a_O_*=0.5, and *R*=0.2*a_O_*. Zoomed-in (grey boxes) drift-diffusion dynamics, occurring on slow time scales. **d,** Comparing deterministic cellular automata (left column) and stochastic cellular automata (right column) with the same parameter values in the activate-deactivate phase (*N*=441, *R*=0.2*a_O_*, *a_O_*=0.5, *K*=10, *C_ON_*=5, and noise strength *ξ* =0.1(for “with noise” automata)). 100 trajectories (red streamlines), each starting at *p*=0.5 and *I* ≈ *0*, shown for both types of automata. Black dots indicate terminal points of each trajectory. Dashed black curve is the theoretically predicted maximum value of *I* as a function of *p* (same as in Fig. 3). Dashed orange curve is the theoretically predicted maximum value of *I* as a function of *p* when noise-driven ordering is present (formula in Supplementary section 4.5). **e,** Average number < *t_final_* > of time steps taken by stochastic cellular automata to terminate as a function of the initial value of *p*, *p_initial_*. Each black data point is an average over at least 1000 cellular automata runs for each value of *p_initial_*. Red curves are analytical estimates based on our theory of drifting-and-diffusing particles with biological noise in phase space.

While a very strong noise can drive a cellular lattice to a “collapse”, in which either all cells turn on or off (Fig. 4b – right column) and a sufficiently low noise keeps a steady state spatial pattern intact (Fig. 4b – left column), we found that a moderately strong noise can drive a steady state spatial pattern to a metastable spatial pattern in which not every cell is on or every cell is off. This is most evident in the activate-deactivate phase (Figs. 4c and 4d; Supplementary Fig. 21). In the activate-deactivate phase, metastable patterns can exhibit glassy dynamics: the *p*, *I*, and *h* can change so slowly that over hundreds of time steps, the spatial pattern essentially remains the same (Fig. 4c). By comparing multiple particle trajectories produced by the noisy cellular automata with those produced by the deterministic cellular automata in the activate-deactivate phase, we found that a moderately strong noise can liberate the trapped particles and let them roll down further in the pseudo-energy landscape (Fig. 4d). These liberated particles increase their values of *I*, often well above the previously determined *I_max_* (Fig. 4d – black curve), until they eventually stop in new traps that are deeper than their previous traps, from which the particles can escape if the noise were stronger (Supplementary section 4.5). But the particles still cannot have an arbitrarily high value of *I* as the *I* is still bounded above when a moderate noise drives the disorder-to-order dynamics (Fig. 4d – orange curve; details in Supplementary section 4.5).

By noting that we can combine the biological noise with the diffusion process that is associated with our ignorance of the microstates, we could extend the equations of motion for the deterministic cellular automata (equations (3) and (4)) to correctly describe the particles drifting and diffusing in the noisy cellular automata (Supplementary section 4.5). Our equations of motion with the biological noise recapitulates the main statistical features of the noisy cellular automata, including the mean halting times <*t_final_*> (Fig. 4e and Supplementary Figs. 22-23).

As our final step, we sought to extend our framework to lattices containing an arbitrary numbers of cell types and signals. We can generally classify cells that communicate through diffusing signals as engaging in either autocrine- or paracrine-signalling^20^ (Fig. 5a). Paracrine signaling requires at least two types of cells whereby one cell secretes but cannot sense the signal while the other cell senses but cannot secrete the signal^20^. In order to treat paracrine signaling, we first asked how the dynamics of our autocrine cells that we have treated so far change if we remove the self-interaction. Without the self-interaction, we found that the deterministic cellular automaton can operate in a fewer phases: the activate-deactivate phase is the only conducting phase and the autonomy phase no longer exists (Supplementary section 5.1). This is because the cells now operate in an open-loop feedback regime in which its state is controlled solely by the other cells. This can cause the spatial pattern to oscillate between two closely related spatial patterns, as if noise existed, without reaching a true steady state (Supplementary Figs. 24 and 25; details in Supplementary section 5.1). Despite the oscillations, we found that the spatial order still increases over time and that the pathway trapping still occurs that allows for stable steady state patterns (Supplementary Fig. 24).

**Figure 5.**
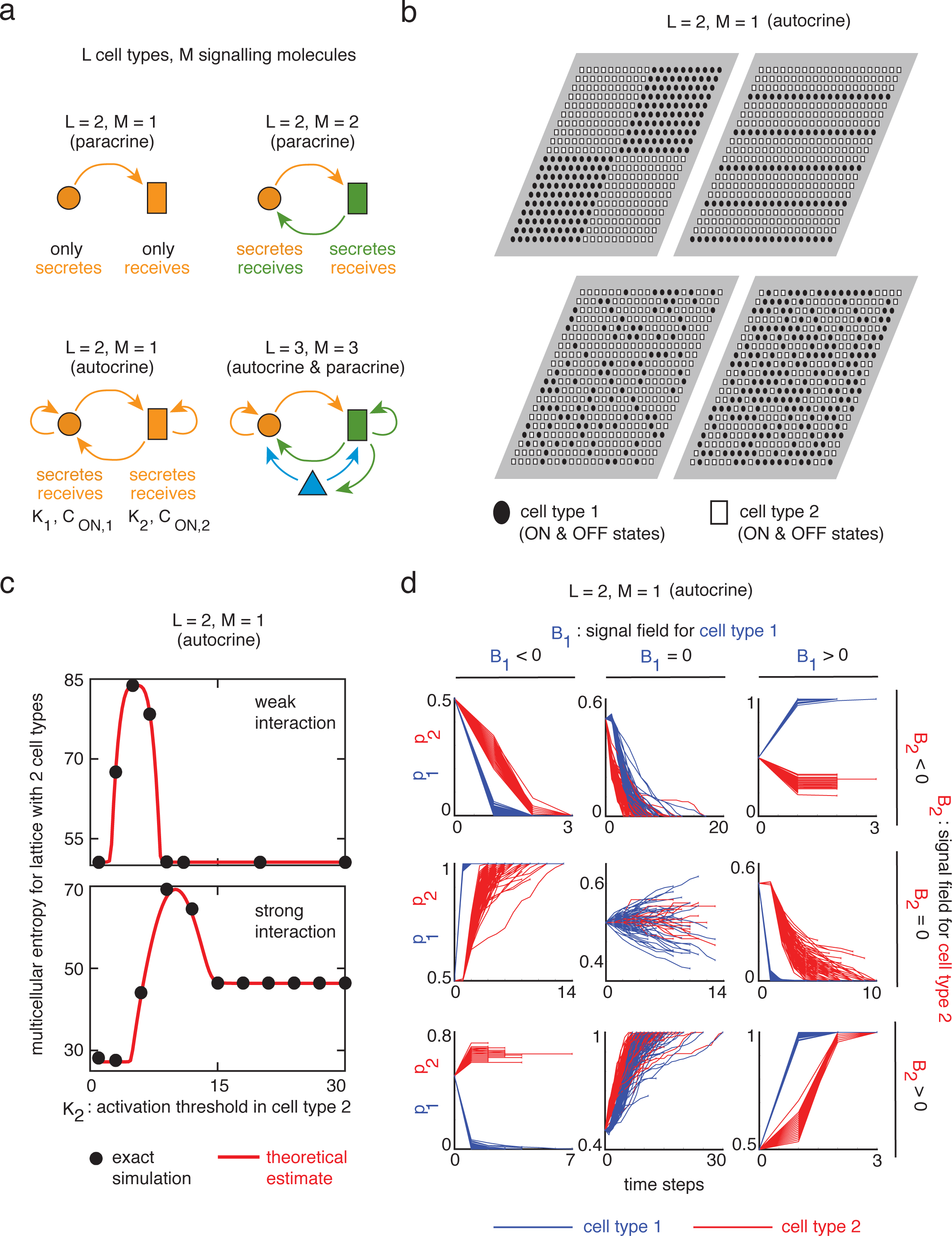
Statistical dynamics of cellular automata with any number of cell types and signalling molecule types. **a,** Examples of autocrine signalling (i.e., a cell secretes a signal and also senses it) and paracrine signalling (i.e., one cell secretes but does not sense the signal while another cell senses but does not secrete that signal) that *L* types of cells with *M* types of signalling molecules can engage in. **b,** Examples of triangular lattices with two types (*L*=2) of autocrine cells (circles and rectangles) that share the same signalling molecule (*M*=1). There are *N* cells in total: *N_1_* cells of type 1 with radii *R_1_* and *N_2_* cells of type 2 with radii *R_2_*. For each lattice, *N*=400, *a_o_*=0.5, *R*=0.2, and *f_N_*=2.358. Using the formalism detailed in Supplementary section 5.2, we obtain interaction strength *f_Im_* between cell type *l* and cell type *m*: (top left) - *N_1_*=200 *N_2_*=200, *f_11_*= 0.854, *f_12_* = 0.325, *f_22_* = 0.854; (top right) - *N_1_*=100, *N_2_*=300, *f_11_* = 0.142, *f_12_* = 0.447, *f_22_* = 1.321; (bottom left) - *N_1_*=100, *N_2_*=300, *f_11_*= 0.147, *f_12_* = 0.443, *f_22_* = 1.326; (bottom right) - *N_1_*=200, *N_2_*=200, *f_11_* = 0.585, *f_12_* = 0.594, *f_22_* = 0.585. **c,** Analytical estimate (red) and exact values from automata (black) for the multicellular entropy of cellular lattices with two autocrine cell types with a shared signal (*L*=2, *M*=1) as in (b). (Top - weak interaction): *a_o_*=1.5, *R_1_*=*R_2_*=0.2*a_o_*, *N_1_*=78, *N_2_*=43, *K_1_*=10, *C_ON,1_*=10, *C_ON_,_2_*=5. (Bottom - strong interaction): *a*_o_=0.5, and others are the same as the top panel. **d,** Fraction *p_1_* of cells of type 1 that are ON (blue curves) and fraction *p_2_* of cells of type 2 that are ON (red curves) obtained through cellular automata for two types of autocrine cells with shared signal (*L*=2, *M*=1, *N*=400, *N_1_*=120, *N_2_*=280, *a_O_*=0.5, *R_1_*=*R_2_*=0.2*a_O_*, *C_ON,1_*=8, and *C_ON,2_*=5). All start with *p_1_*=*p_2_*=0.5. *B_u_* is the signal field for cell type *u* (*u*=1, 2). The signs of *B_u_* determine cell type k’s change in *p_u_* over time. If neither *B_1_* nor *B_2_* is zero, then *p_u_* increases over time if *B_u_* > 0 and decreases over time if *B_u_* < 0. When *B_u_* ≈ 0, then *p_u_* nearly stays the same for the first several time steps and then follows the direction of change of the other cell type’s *p*.

To fully treat cellular lattices composed of arbitrary numbers of cell types and of signal types (including multiple autocrine and paracrine signals) (Fig. 5b), we now allow for each cell type *m* to have a distinct genetic circuit with an activation threshold *K_m_*, signal concentration *C_ON_*,*_m_* that a lone ON-cell generates on itself, and a signal concentration *C_OFF_*,*_m_* that a lone OFF-cell generates on itself. We also generalize the interaction strength *f_N_*(*a_o_*) to an interaction strength matrix *F* whose submatrix *F_mn_* with elements *f_ij_* quantifies how cell-i of type *m* interacts with cell-j of type *n* (examples in Fig. 5b; details in Supplementary section 5.2). For instance, if the cell type *m* only engages in paracrine signalling or there is no cross-signalling between cells of types *m* and *n*, then *F_mm_*=0. As an example of this approach, we extended our framework to define and obtain an explicit formula for the multicellular entropy *S* = *log*(*Ω_E_*) for lattices with two cell types and one autocrine signal. This formula closely agrees with the exact values of *S* obtained from the cellular automata (Fig. 5c). Our approach also enabled us to extend the multicellular Hamiltonian *H* to lattices of *L* cell types:

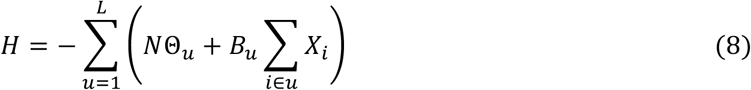

where the first sum runs over all cell types and the internal sum runs over all cells of the same type. In equation (8), we have defined the normalized spatial order of cell type *u* as 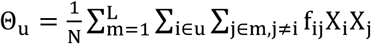 and the signal field for cells of type *u* as 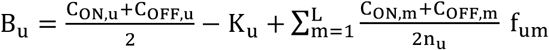, where *n_u_* is the fraction of cells that are of type *u* and 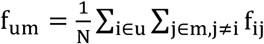. As in the case of lattices composed of identical autocrine cells, we found that this Hamiltonian also monotonically decreases over time (details in Supplementary section 5.3). Also, as in the lattices of identical autocrine cells, the signs of the signal fields *B_u_* help us determine how the fraction *p_u_* of cells of type *u* that are ON changes over time (Fig. 5d). Namely, when *B_u_* is not zero for any cell type, then *p_u_* increases over time whenever *B_u_* > 0 (i.e., activation of cell type *u*) and decreases over time whenever *B_u_* < 0 (i.e., deactivation of cell type *u*). When the signal field *B_u_* for cell type *u* is zero but the signal field *B_q_* for any cell type *q* is non-zero, *p_u_* can remain nearly unchanged for the first few time steps while *p_q_* changes, and then *p_u_* sharply increases (if cell type *q* is activated) or decreases (if cell type *q* is deactivated) due to the increasing influence of the cells of type *q* on the cells of type *u* (Fig. 5d). In all cases, we can show that the spatial order increases over time (details in Supplementary section 5.3). When the signal field for all cell types is zero, highly organised metastable patterns can again occur due to pathway trapping. In summary, we found that a cellular lattice consisting of *L* cell types acts as *L* distinct particles that drift-and-diffuse in a common phase space spanned by *p* and *I*, with the location of particle-u being (*p_u_*, *I_u_*). Specifically, the *L* particles interact among them. This yields a pseudo interaction potential (denoted by the first term in equation (8)). Moreover, each particle interacts with its own signal field. This yields a single-particle pseudo-energy (denoted by the second term in equation (8)). The resulting collective motion of the *L* particles in the phase causes the total pseudo-energy *H* (equation (8)) to monotonically decrease over time.

Here we have presented the core principles that drive the disorder-to-order dynamics in a general class of cellular automata. The cellular automata that we have investigated here are formalized idealizations, based on experimental findings, of spatial pattern forming multicellular systems such as tissues (Supplementary Table 1 and Supplementary section 1.1). Our statistical mechanics-type framework allows one to analytically investigate a class of cellular automata that have largely been studied through exhaustive simulations. Specifically, we have shown that for this class of cellular automata, which has different operational phases (Fig. 1b), we can group cellular lattices composed of a single type of autocrine cell with a positive feedback into macrostates defined by the fraction *p* of cells that are ON and the spatial order parameter *I*. We have shown that the particle’s position (*p*, *I*) in the phase space evolves over time according to Langevin dynamics – the particle drifts and diffuses according to probability distributions that depend on the strength of non-local interactions among cells and the parameters of the genetic circuits. We have shown that underlying these drift-and-diffuse dynamics of particles is a pseudo-energy landscape, whose “altitude” is defined by a multicellular Hamiltonian, which is a function of *p* and *I* and shares similarities with, along with stark differences from, the Hamiltonians of spin glasses and the Hopfield network. Moreover, we found that the interdependence of *p* and *I* and geometric constraints on the lattice lead to pathway trapping – the trapping of the particles along inclined regions of the pseudo-energy landscape that causes metastable spatial patterns. We have shown that adding noise to each cell in the cellular automata yields metastable spatial patterns whose spatial order is much higher than those generated without noise and whose stability eventually breaks down due to slowly varying changes that are reminiscent of glassy dynamics. Finally, we have shown that our framework could be extended to cellular automata with an arbitrary numbers of cell types and of signal types.

Previous studies have provided deep insights into how efficiently genetic networks in a cell can process information to generate accurate input-output relationships in which the input usually is a steady state gradient of signal that a cell detects and the output is the cell’s steady state gene expression^12-19^. For example, researchers have quantified the amount of information encoded in gene expressions of fly embryos^19^ and mouse cells^15^. Linking such studies – which deal with the performance efficiencies of genetic circuits based on the steady or quasi-steady state gene expression levels^12,14,19^ – with our mechanistic description of multicellular dynamics – including changes in gene expression over time, dynamics of signal generation and sensing, and changes in spatial patterns over time – may provide a way forward to formally address questions regarding spatiotemporal dynamics of information processing that have been underexplored. For example, it may be possible to define the number of bits of information encoded in an initial spatial pattern and then determine how a cellular automaton propagates this information over time due to the signals shared among the cells to eventually arrive at a stable spatial pattern.

Another underexplored topic that our work may provide a way to address is the idea that we may formally view biological processes as cells solving problems by using, potentially yet-to-be identified, computational algorithms^24,29,30^. For example, studying the collective dynamics of the sensory organ precursor cells in the fly *D. melanogaster* led to a more efficient computational algorithm^24^. Conversely, researchers have shown that robots, like cellular automata, that follow a small set of rules to interact with each other can recapitulate the collective behaviours of organisms such as termites^25^. But it has been difficult to generalize these observations to advance the idea that computational algorithms are implemented by networks of cells and genes due to a lack of a formalized framework to analyze the dynamics of large, entangled cellular networks. Future studies may resolve this challenge by using the macrostate functions that we presented here to deduce the statistical dynamics of cellular automata that follow rules other than the ones that we have investigated here. Moreover, our study could be extended to determine the computational complexity of algorithms that networks of cells use. For example, we may view the termination time *t*_final_ of the cellular automaton (Fig. 2d) as the run time of an underlying algorithm that cells use to complete a certain task. Then we can determine the cellular algorithm’s computational complexity by finding how *t_final_* scales as a function of the size of the “input” that the cells use (e.g., as a function of the number of cells). Viewing biological processes as formalized computational algorithms in which signals are transmitted and processed over space and time as in cellular automata^29^, may reveal central principles that govern signal-processing dynamics of biological networks. Moreover, since the cellular automata of the type that we studied here can represent non-living systems such as networks of signal-relaying switches that interact through diffusing signals, our approach may aid in better understanding network-level dynamics of signal-processing physical systems.

## METHODS

### Multicellular entropy

A full derivation is in Supplementary section 1.3. We treat the state of each of the *N* cells as a random variable, with the constraint that *p* is fixed. If we consider only the signal that a cell senses due to the secretion of all the other cells (i.e., excluding the signal secreted by the cell itself), its concentration 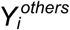 obeys the following statistics

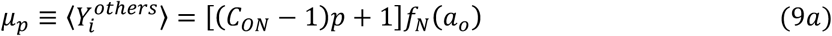

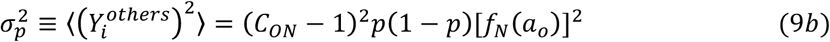

where *µ_n_* and *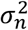* are mean and variance of 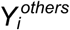 respectively that are determined by averaging over all the 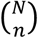 spatial patterns that have *n* ON-cells. By assuming that the probability *P_ON-ON_* that an ON-cell stays on and the probability *P_OFF-OFF_* that an OFF-cell stays off are normally distributed, it follows that *µ_n_* and 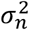 specify the fraction of the 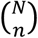 spatial patterns that are steady states. Summing over all *n* yields an estimate for *S* that closely matches the exact value of *S* obtained from running the cellular automaton (Fig. 1d). The total number Ω_E_ of steady state spatial patterns that a cellular lattice can have is the sum of the number Ω_p_ of steady state patterns for each value of *p*,

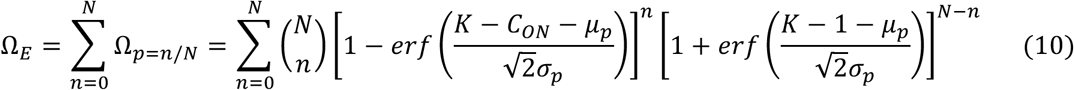

This result agrees well with the exact value obtained from the cellular automaton runs (Fig. 1d).

### Transition matrix formalism

Full details are in Supplementary section 2.3. Consider a randomly picked spatial pattern with *n* ON-cells. The transition matrix Ξ determines the probability that one spatial pattern becomes another spatial pattern after one time step, for any pairs of spatial patterns and yields a macrostate-level description. Specifically, its matrix element 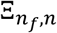, is the probability that, if we randomly pick one out of the 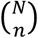 spatial patterns with *n* ON-cells, the selected pattern will have *n_f_* ON-cells in the next time step. Using the probabilities *P_ON-ON_* and *P_OFF-OFF_* that we computed for the multicellular entropy, we can calculate the probability that a cell will change its state, which is *P_ON-OFF_ = 1 - P_ON-ON_* for ON-cells and *P_OFF-ON_ = 1 - P_OFF-OFF_* for OFF-cells. We then use these to calculate the probability that *y*_+_ cells will turn on and *y*_-_ cells will turn off in the next time step. Summing the probabilities *p*(y_-_, y_+_; n) for all possible pairs (*y*_+_, *y*_-_), we obtain the transition probability 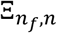.

### Mean-field approximation

Full details are in the Supplementary information (section 3.1 and Appendix A). With our periodic boundary condition (see Supplementary section 1.1), the signal concentration 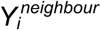 on cell-i due to everyone else is 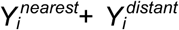, where 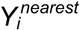 is due to its nearest neighbours and 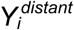 is due to all the other cells. For a mean-field approximation, we calculate 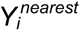 exactly by using the number 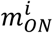 of the nearest neighbours of cell-i that are ON and 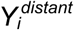. Using this mean-field approach, which ignores the precise location ON/OFF-cells other than those that are the nearest neighbours, we can compute *p*. Moreover, using this approach, we can show that the average number of ON-neighbours < *m_ON_* > monotonically increases over time.

### Langevin dynamics in the phase space spanned by *p* and *I*

Full details are in Supplementary section 2.4. Equation (3) describes the particle (cellular lattice) dynamics, in terms of the macrostate variables *p* and *I*, with the Langevin approach. This description requires calculating the drift and the noise for both *p* and Θ. The probability that an ON-cell turn off in the next time step is 1 − *P_ON–ON_*. The probability that an OFF-cell turns on in the next time step is 1 − *P_OFF–OFF_*. Then if we assume that *N* is sufficiently large so that *p* assumed to be a continuous variable, we have

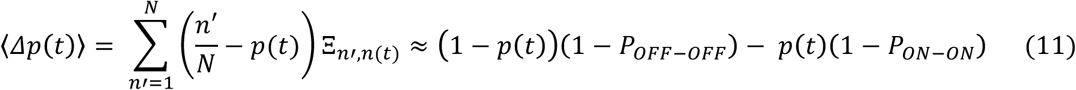

Analogously, we can estimate the variance of the noise *η*_Δ*p*_ by assuming that the switching of the ON- and OFF-cells are independent of each other:

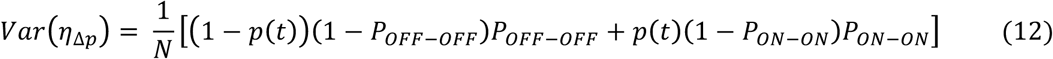

For the normalized spatial order parameter 0, we can use its definition deduce that 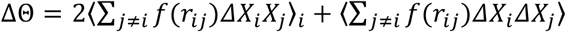 and then calculate the mean and the variance of both terms in this expression. We approximate the second term in the expression as 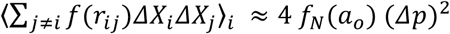 and use the mean and the variance calculated for Δ*p* to proceed further. For the first term in the expression for ΔΘ, we have

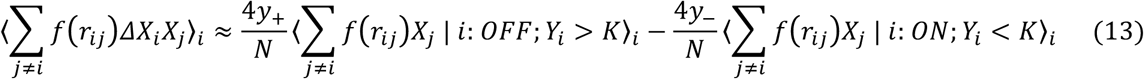

Invoking the Central Limit Theorem, both averages in the above expression (averaged over all cells) can be thought of as truncated Gaussian distributions from which the mean and variance are calculated to determine 〈*ΔΘ*(*t*)〉 and *Var*(*η*_Δ*Θ*_).

### Proof that the multicellular Hamiltonian monotonically decreases over time

Full details are in Supplementary section 3.3. Here we outline the proof. In the multicellular Hamiltonian (equation (6)), we have three terms to consider. The third term is a constant. The first term monotonically decreases over time because < *m_0N_* > monotonically increases over time, as we have shown with the mean-field approach (Supplementary section 3.1). The second term in equation (6) is negative when 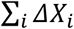 and *B* have opposite signs. For the cellular automaton that operate in the insulating phases, *ΔX_i_* = 0 and consequently *ΔH* = 0. In the activate phase, we have *B*>*0* and in the deactivate phase, we have *B*<*0*, implying that the second term decreases in those phases. In the activate-deactivate phase, there is activation when *B*>*0* and deactivation when *B*<*0* (Supplementary Figure 18). Putting everything together, we conclude that the multicellular Hamiltonian monotonically decreases over time.

### Monte Carlo simulation of statistical dynamics based on equations of motion (equation 3)

Full details are in Supplementary section 2.5. We performed the Monte Carlo simulations of the drifting-and-diffusing particles with the Euler Method for stochastic differential equations. Based on the current macrostate (*p*(*t*), *Θ*(*t*)) (i.e., particle’s current position), we calculate a drift (〈Δ*p*(*t*)〉,〈*ΔΘ*(*t*)〉) and sample two random numbers using the Marsaglia polar method with variance *Var*(*η_Δp_*) and *Var*(*η_ΔΘ_*) to calculate the subsequent macrostate (*p*(*t* + l), *Θ*(*t* + 1)). This macrostate has a probability 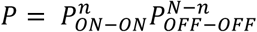 of being a steady state. Therefore, we uniformly sample a number *z* between 0 and 1. If *z* < *P*, then we consider the spatial pattern to be in a steady state. Running this procedure several times, it is possible to realize many different particle trajectories, which we compare with those obtained by the cellular automaton (Figures 2c and 2d).

### Extension of our formalism to multiple cell types with multiple signals

Full details are in Supplementary section 5.2. For a lattice with multiple cell types, we replace the scalars *C_ON_, C_OFF_* and *K* by vectors that are ordered by the cell type. For instance, if we have *L* cell types, we have 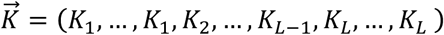. Moreover, the interaction terms *f*(*r_ij_*) for an interaction matrix *F* = ⟦*F_lm_*⟧ where *F_lm_* contains all the interactions terms *f*(*r_i_*_∈*l*, *j*∈*m*_) among cells of type *l* and cells of type *m*. With this notation, we can extend the multicellular Hamiltonian to lattices with an arbitrary number of cell types (equation (8)). Moreover, we can extend our calculations of *P_ON–ON_* and *P_OFF–OFF_* to obtain probabilities *P_ON–ON;l_* and *P_OFF–OFF;l_* that a cell of type *I* remains in its ON and OFF states, respectively, in the next time step. The multicellular entropy is then given by

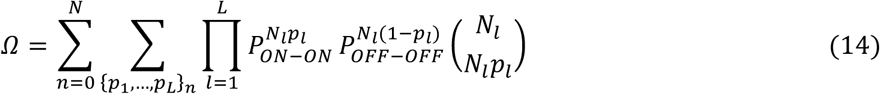

where *N_l_* is the number of cells of type *I*, *p_l_* is the fraction of cells of type *l* that are ON, and {*p*_1_, …, *p_L_*}*_n_* represents all the possible combinations of *p_I_*’s with the constraint that there is a total of *n* ON-cells on the lattice. This result closely agrees with the exact values obtained for lattices with two cell types (Figure 5c).

## ADDITIONAL INFORMATION

Full details of derivations, in-depth discussions of the results summarized here, supplementary text, and supplementary figures are available in the Supplementary Information.

## ACKNOWLEDGMENTS

We thank Y. Blanter (Quantum Nanoscience dept., TU Delft), L. Reese (Bionanoscience dept., TU Delft), A. Raj (Bioengineering dept., Univ. of Pennsylvania) and the members of the Youk group for insightful discussions and comments on the manuscript. HY is supported by the European Research Council’s Starting Grant (#677972), the Netherlands Organization for Scientific Research (NWO) Vidi award (#680-47-544), and the NWO NanoFront program.

